# GeneRax: A tool for species tree-aware maximum likelihood based gene family tree inference under gene duplication, transfer, and loss

**DOI:** 10.1101/779066

**Authors:** Benoit Morel, Alexey M. Kozlov, Alexandros Stamatakis, Gergely J. Szöllősi

## Abstract

Inferring phylogenetic trees for individual homologous gene families is difficult because alignments are often too short, and thus contain insufficient signal, while substitution models inevitably fail to capture the complexity of the evolutionary processes. To overcome these challenges species tree-aware methods also leverage information from a putative species tree. However, only few methods are available that implement a full likelihood framework or account for horizontal gene transfers. Furthermore, these methods often require expensive data pre-processing (e.g., computing bootstrap trees), and rely on approximations and heuristics that limit the degree of tree space exploration. Here we present GeneRax, the first maximum likelihood species tree-aware phylogenetic inference software. It simultaneously accounts for substitutions at the sequence level as well as gene level events, such as duplication, transfer, and loss relying on established maximum likelihood optimization algorithms. GeneRax can infer rooted phylogenetic trees for multiple gene families, directly from the per-gene sequence alignments and a rooted, yet undated, species tree. We show that compared to competing tools, on simulated data GeneRax infers trees that are the closest to the true tree in 90% of the simulations in terms of relative Robinson-Foulds distance. On empirical datasets, GeneRax is the fastest among all tested methods when starting from aligned sequences, and it infers trees with the highest likelihood score, based on our model. GeneRax completed tree inferences and reconciliations for 1099 Cyanobacteria families in eight minutes on 512 CPU cores. Thus, its parallelization scheme enables large-scale analyses. GeneRax is available under GNU GPL at https://github.com/BenoitMorel/GeneRax.

## Introduction

Reconstructing the evolutionary history of homologous genes constitutes a fundamental problem in phylogenetics, as the phylogenetic trees of gene families (henceforth, gene family trees) play a prominent role in numerous biological studies. For instance, gene family trees (GFTs) are essential to understand genome dynamics (Touchon *et al*., 2009), to study specific traits (Musilova *et al*., 2019), or to infer the species tree (Boussau *et al*., 2012; Mirarab *et al*., 2014).

Standard phylogenetic methods infer trees from multiple sequence alignments (MSAs), for instance using the maximum likelihood (ML) criterion (Kozlov *et al*., 2019; Nguyen *et al*., 2015). Under the correct substitution model, ML methods are statistically consistent (Yang, 1994), that is, they converge to the true tree when the sequences are long enough. However, this condition is often violated for GFTs: typical pergene MSAs are short (50 to 1000 sites) and can comprise a large number of sequences representing a large number of *taxa* (hundreds or thousands for large gene families). As a result, there is typically insufficient signal in the MSA to reconstruct a well supported phylogeny. In other words, the tree with the highest likelihood might not correspond to the true tree.

Species-tree-aware (STA) approaches aim to compensate for this insufficient phylogenetic signal by relying on a putative species tree. Indeed, GFTs and the species tree exhibit an intricate relationship: genes evolve within a (species) genome and undergo biological processes such as duplication, horizontal gene transfer (HGT), loss, or speciation (Fig. 1). Therefore, although GFTs can be incongruent with the species tree, their own evolutionary history is, to a substantial degree, determined by the species tree. STA methods exploit this relationship between the GFTs and the species tree to leverage additional information for GFT inference. In the following, we denote gene duplication, gene loss, and HGT events as *DTL events*.

**FIG. 1.**
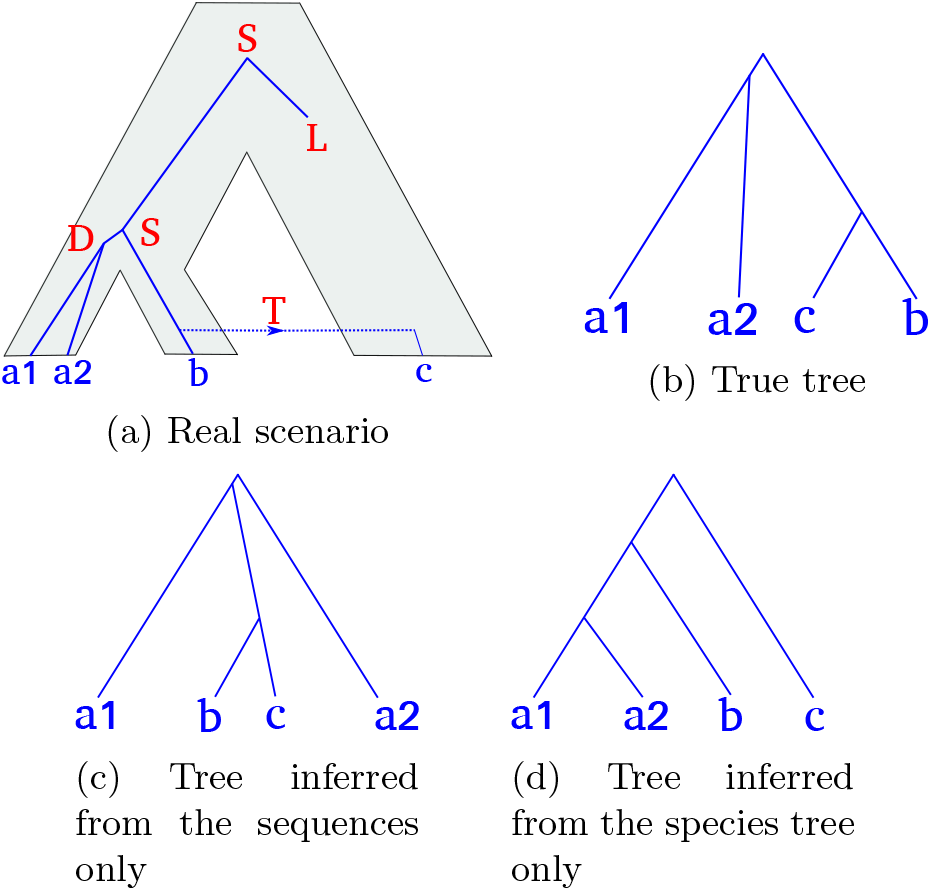
A gene tree evolving along the species tree, and several possible inferred trees. (a) The true history. The gene tree (blue lines) evolves within the species tree (grey area), and undergoes speciations (S), duplications (D), losses (L) and HGT (T). (b) The true gene tree. (c) A gene tree inferred with a sequence-aware method. The duplication and the speciation between the species a and b are very close in time, and there is not enough signal in the sequences to correctly decide which split happened first. (d) Tree inferred from the species tree only (without accounting for the sequences), assuming that HGT are less likely than duplications.

A common approach used by STA methods (Chen *et al*., 2000; Noutahi *et al*., 2016; Scornavacca *et al*., 2014) consists of contracting weakly supported GFT branches into polytomies, which are subsequently resolved using the species tree. These heuristics limit the set of GFTs explored to trees that can be obtained as combinations of alternative resolutions of the contracted branches. Most existing implementations (Chen *et al*., 2000; Noutahi *et al*., 2016) are based on parsimony, and require a priori specification of arbitrary DTL parsimony costs. This is particularly problematic if the substitution model is misspecified, or if it fails fails to adequately capture the complexity of the data. This is commonly the case for shorter gene alignments where parameter rich substitution models are more difficult to use. In addition, the user must define what a “low support value” for branch contraction is, often by setting an arbitrary threshold. Treerecs (Comte *et al*., 2018) addresses this last limitation by exploring several thresholds, and returning the GFT that maximizes a likelihood score that is based on both, the MSAs, and the species tree. Finally, obtaining branch support values usually requires a substantial amount of computational effort (e.g., 1-2 orders of magnitude more than for a simple ML tree search on the original MSA, if the classic Felsenstein Bootstrap is used (Felsenstein, 1985)).

Other STA methods utilize a hierarchical probabilistic model of sequence level substitutions and gene level events, such as duplication, transfer, and loss. This allows the definition of the *joint likelihood* as the product of the probability of observing the alignments given the GFTs (*phylogenetic likelihood*) and the probability of observing the GFTs given the species tree (*reconciliation likelihood*):

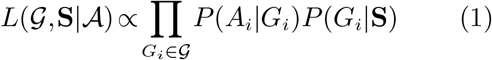

where **S** is the species tree, 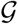 is the set of GFTs, and 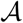 the set of corresponding MSAs. Phyldog (Boussau *et al*., 2012) co-estimates the GFTs and the species tree by conducting a tree search that is based on such a joint likelihood score. However, Phyldog does not model HGT. ALE (Szöllősi *et al*., 2013a) calculates the joint likelihood using a dynamic programming scheme that requires the phylogenetic likelihood to be approximated via conditional clade probabilities (Larget, 2013). In order to calculate conditional clade probabilities, ALE requires a sample of GFTs as input that are typically obtained via Markov Chain Monte Carlo (MCMC) sampling. This approach has two shortcomings. First, the conditional clade probability approximation inevitably limits the set of GFTs explored to trees that are comprised of clades observed in an tree sample, as the phylogenetic likelihood of all other trees is approximated to be zero (Szöllősi *et al*., 2013a). While being less restrictive, conceptually this limitation is nonetheless analogous to those induced by the branch contraction methods discussed above. It is also similarly sensitive to model miss-specification and inadequacy. Secondly, obtaining a tree sample, either via Bayesian phylogenetic MCMC methods or via bootstrap methods for a set of gene families is computationally expensive. For an in depth review of GFT inference methods, see (El-Mabrouk and Noutahi, 2019; Szöllősi *et al*., 2014).

Probabilistic frameworks to model both, sequence (Felsenstein, 1981), and gene evolution events (Åkerborg *et al*., 2009; Sennblad and Lagergren, 2009; Szöllősi *et al*., 2013b) can be found in the literature. However, no ML tool can cutrrently directly infer GFTs from MSAs by simultaneously accounting for sequence substitutions and DTL events. We believe that such a method can substantially improve the accuracy of GFT inference. A common argument against using STA ML approaches is the amount of time and computational resources required to conduct such analyses (El-Mabrouk and Noutahi, 2019). However, a joint (phylogenetic and reconciliation likelihood) ML approach does not require expensive pre-processing and can therefore decrease the overall computational cost substantially, while increasing accuracy at the same time. Tree search heuristics are widely used to infer phylogenies from sequence data (Kozlov *et al*., 2019; Nguyen *et al*., 2015) using the phylogenetic likelihood. Thus, extending these methods by joint likelihood calculations represents a natural way of improving the accuracy of GFT inference.

Here we introduce GeneRax, our novel software to infer ML reconciled GFTs based on a joint reconciliation and phylogenetic likelihood. We use the term *reconciled GFT* to designate both, the GFT topology, and its reconciliation with the species tree. The input for GeneRax consists of a rooted, but undated binary (fully bifurcating) species tree, a set of per-family MSAs (DNA or amino-acid), and corresponding gene-to-species leaf name mappings. Several genes from the same gene family can be mapped to the same species. In addition, the user can provide initial GFTs, typically inferred via standard phylogenetic methods (Kozlov *et al*., 2019; Nguyen *et al*., 2015). GeneRax is easy to use, models DTL events, and can process gene families in parallel. Employing a hierarchical probabilistic model allows it to simultaneously account for both, the signal from the gene family MSAs, and from the species tree. It estimates all substitution and DTL events intensity parameters, and does not require any *ad hoc* threshold nor any arbitrary DTL event parsimony costs.

Nonetheless, one should keep in mind that incomplete lineage sorting (ILS) constitutes another important source of discordance between GFTs and the species tree. A recent study suggests that ILS can bias reconciliation inference (Zheng and Zhang, 2014). To this end, we also assess the impact of ILS on the reconstruction accuracy of STA methods and discuss the limitations of GeneRax in the presence of ILS.

## New Approaches

In this section, we outline the joint likelihood computation, our tree search algorithm, and our parallelization scheme.

### Reconciliation likelihood

In this subsection, we derive the reconciliation likelihood for a rooted GFT given an undated, yet rooted species tree, as implemented in ALE.

The “undated” DTL model, in contrast to the continuous time model described in (Szöllősi *et al*., 2013b), is a discrete state model, which starts with a single gene copy on a branch of the species tree. Subsequently, gene copies evolve independently until, either all copies are observed at the leaves, or every gene copy becomes extinct. On an arbitrary branch of the species tree a gene copy:

- either duplicates and is replaced by two corresponding gene copies on the same branch (with probability *p^D^*)
- a new copy is transferred to a random branch that is *not* ancestral to the donor branch, but otherwise drawn uniformly at random from the species tree, while a copy also remains on the donor branch (with probability *p^T^*)
- is lost (with probability *p^L^*)
- undergoes a speciation event on internal branches, in which case it is replaced by a copy on each descendant branch (with probability *p^S^* = 1 − *p^D^* − *p^T^* − *p^L^*)
- is observed for terminal branches, that is, arrives in the present and is observed, thus terminating the process (again with probability *p^S^* = 1 − *p^D^* − *p^T^* − *p^L^*)

By *δ*, λ, and *τ* we denote the duplication, loss, and transfer intensity parameters that parametrize the above event probabilities as follows:

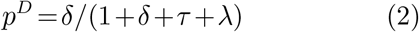

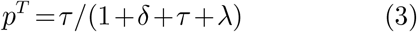

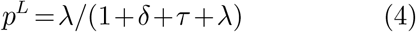

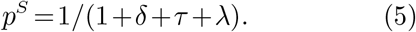

The probability of observing a rooted GFT **G** under the undated DTL model defined above can be calculated by summing over all possible series of D, T, L, and S events (henceforth “scenarios”) that yield a rooted topology that is congruent with **G**. The sum over all possible scenarios is computed in two steps (Sjöstrand *et al*., 2013; Szöllősi *et al*., 2013b). First, we calculate the extinction probability of a gene copy that was initially present on some branch of the species tree. The extinction probability is the sum over all scenarios that do not yield descendants. Second, we sum over all reconciliations of **G**, where a reconciliation of **G** corresponds to a specific sequence of D, T, S, and gene copy extinction events, and its probability corresponds to the product of the specific sequence of events (cf. Fig. 2a).

**FIG. 2.**
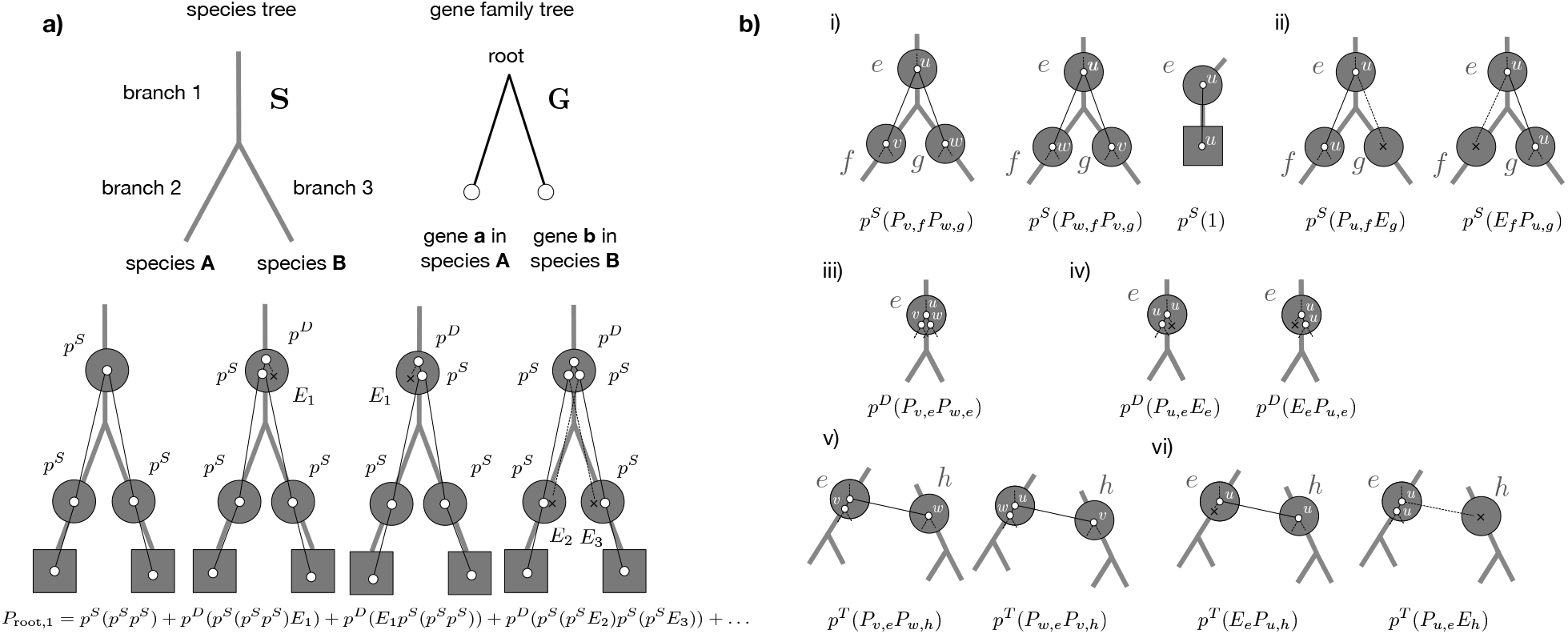
Calculating the probability of G along S. a) The probability that the rooted GFT **G** is generated along the rooted species tree **S** according to the “undated” DTL process can be calculated by summing over all reconciliations. Here we show the leading terms in the sum over all reconciliations that start with a single gene copy on branch 1 of **S** and obtain a rooted gene tree that is congruent with **G**. b) More generally, to calculate *P_u,e_*, that is, the sum over all reconciliations generating the sub-tree below some internal node *u* of **G** starting from a single gene present on the internal branch *e* of **S** we must consider the following events i) if *e* is an internal branch of **S**, speciation with probability *p^S^* such that the descendants of *v* on **G** are observed on *f* of **S** and of *w* on **G** are observed on *g* of **S**, or vice versa. If *e* is a terminal branch, with probability *p^S^* gene *u* will be observed at the terminal branch *e*; ii) if *e* is an internal branch of **S**, speciation with probability *p^S^* such that the descendants of *u* are observed on *f* of **S** and the copy on *g* goes extinct with probability *E_g_*, or vice versa; iii) duplication with probability *p^D^* such that *v* and *w* are both observed on *e*; iv) duplication with probability *p^D^* such that either the first or second copy goes extinct, each with probability *E_e_* and *u* is observed on *e*; v) transfer with probability *p^T^*, such that the respective branches *v* and *w* correspond to the copy on the donor branch *e* of **S**, while the other copy corresponds to the recipient copy on branch *h* of **S** that is not an ancestor of *e* and finally vi) transfer with probability *p^T^* followed by the extinction of either, the copy in the donor linage *e* with probability *E_e_* of the extinction of the copy in the recipient, with probability *E_h_*. These correspond to the terms of Eq. 2.

To begin, let *e* be branch of the species tree **S**, and let *f* and *g* be its descendant branches (remember that the species tree is rooted). Let 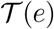 be the set of species tree branches that can receive a HGT from *e*. Because we do not assume any time information on the species tree other than the order of descent induced by the rooted tree topology, we consider that 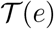 corresponds to all nodes that are not ancestors of *e*. We allow transfers from *e* to its descendants, because a gene could have evolved along an extinct or unsampled lineage and could subsequently have been transferred back to a descendant of *e* (Szöllősi *et al*., 2013b).

The extinction probability, that is, the probability that a gene copy observed on an internal branch *e* becomes extinct before being observed at the tips of the specie tree is:

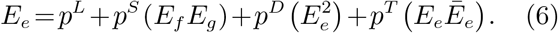

The terms correspond to the i) loss probability, ii) speciation and subsequent extinction probability in both descending lineages (this term must be omitted for terminal branches), iii) duplication and subsequent extinction probability of both copies and finally iv) transfer and subsequent extinction probability of both, the donor copy on branch *e*, and the transferred copy on branch *h*. For the latter event we have introduced the notation:

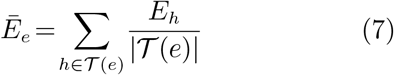

In (6), the value of *E_e_* depends on *Ē_e_*, and thus on the extinction probabilities of all species in the species tree. We iteratively estimate *Ē_e_* and *E_e_* for all nodes *e* in the species tree, by initializing [*E_e_*]^0^ = 0 and computing:

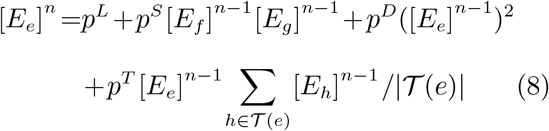

If the limit of the sequence [*E_e_*]^*n*^ exists, then it represents the solution of (6). We do not prov the existence of this limit here.

In simulations, we observed that 5 iterations are sufficient to estimate *E_e_*, and we have thus set the number of iterations to 5 in our implementation. In the special case where *τ* = 0 (no HGT), the contribution of the term *Ē_e_* zero, and we can directly compute *E_e_* from *E_f_* and *E_g_*.

To calculate the probability of a rooted GFT **G**, we have to sum over all reconciliations of **G**. This includes D, T, S, and gene extinction events that may have generated the observed, rooted GFT along the species tree. The algorithm to calculate the sum over all reconciliation histories proceeds from the tips of the rooted species tree and rooted GFT toward their respective roots. Let *v* and *w* be descendants of *u* on **G**, and *f* as well as *g* be descendants of *e* on the species tree **S**. For calculating the recursive sum over reconciliations, consider *P_e,u_*, as the sum over all reconciliations that generate the sub-tree below some internal node *u* of **G** starting from a single gene being present on the internal branch *e* of the species tree **S**. We calculate *P_e,u_* by enumerating all possible single **D, T**, and **S** events that can result from *u* on *e*. These are shown in Fig. 2, and yield:

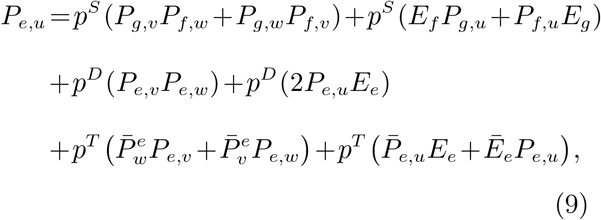

where we have introduced the notation:

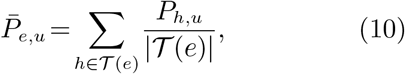

where 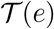 denotes the branches of **S** that are *not* ancestors of *e*.

Similar to the expression for the extinction probability, *P_e,u_* depends on itself. We solve this through fixed point iteration analogously to (6). Aside of the self dependence, every other term involves either descendant branches in **G** (*u* and *w*), descendant branches in **S** (*f* and *g*), or both. This allows to devise a bottom-up dynamic programming recursion starting at the leaves, such that for leaf *g* of the GFT and leaf *s* of the species tree *P*(*g,s*) = 1, if gene *g* maps to species *s*, and zero otherwise.

Given the above, to calculate the reconciliation likelihood, let **G** be a rooted GFT, *r* its root, **S** a rooted species tree, *V*(**S**) the set of nodes of **S**, and *N* = {*δ*, *τ*, λ} the set of DTL intensity parameters. The reconciliation likelihood can then be expressed as:

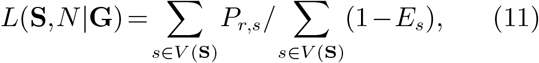

where we divide by 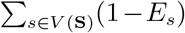 to condition on survival, as extinct gene families cannot be observed.

### Joint likelihood evaluation

GeneRax attempts to maximize the joint likelihood defined as:

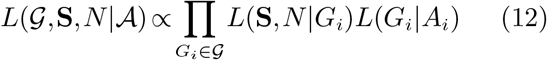

where 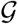 is the set of GFTs, **S** is the species tree, *N* are the DTL event intensity parameters, and 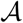 is the set of gene feamily MSAs.

GeneRax estimates the reconciliation likelihood *L*(**S**,*N*|*G_i_*) based on the dynamics programming recursion described above. It uses the highly optimized *pll-modules* library (Darriba *et al*., 2019) to compute the phylogenetic likelihood *L*(*G_i_*|*A_i_*). Hence, GeneRax offers all substitution models also supported by RAxML-NG (Kozlov *et al*., 2019).

### Joint likelihood optimization

Given a set of MSAs and a species tree, GeneRax searches for the set of rooted GFTs and DTL intensity parameters that maximizes the joint likelihood. We illustrate the search procedure in Fig. 3.

**FIG. 3.**
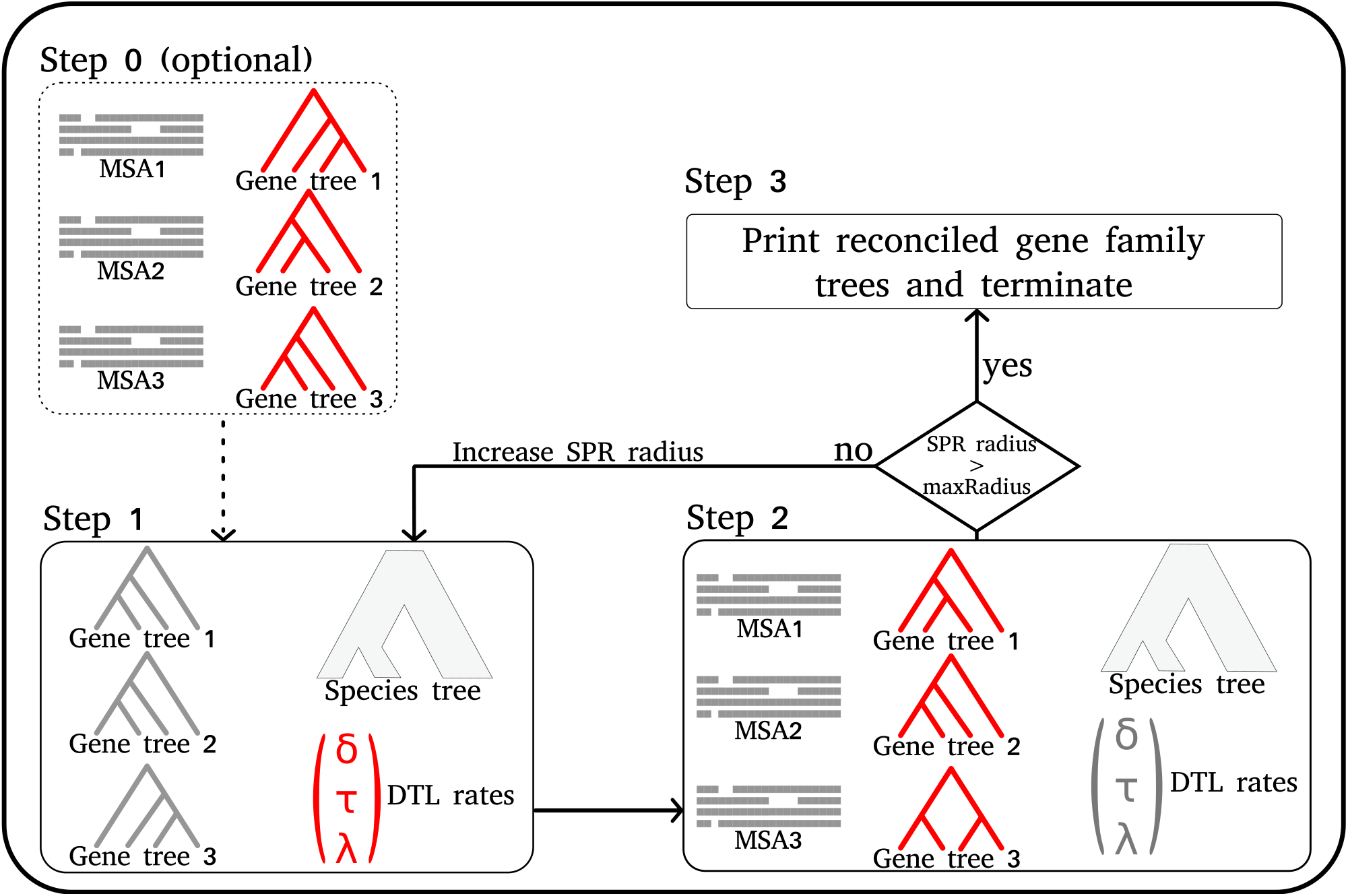
GeneRax pipeline. In each step, we draw in red the parameters that GeneRax optimizes, and in grey the fixed parameters that GeneRax uses to compute the likelihoods. GeneRax performs Step 0 only when starting from random GFTs, to infer ML GFTs from the MSAs. Step 1 optimizes the DTL event rates from the GFTs and the species tree. Step 2 optimizes the GFTs from the MSAs, the species tree and the DTL rates. GeneRax repeats Step 1 and Step 2 with increasing SPR radius, until it reaches the maximum radius. Then it applies Step 3 to reconcile the GFTs with the species tree.

GeneRax either starts from user-specified GFTs or from random GFTs. Our joint likelihood search algorithm needs to start from GFTs with high phylogenetic likelihood, preferably inferred with phylogenetic ML tools such as RAxML-NG (Kozlov *et al*., 2019). We provide a rationale for this in the Results section. When starting from random GFTs, GeneRax performs an initial search (Step 0 in Fig. 3) that solely maximizes the phylogenetic likelihood, without accounting for the reconciliation likelihood.

After this optional step, GeneRax starts optimizing the joint likelihood, by alternating between optimizing the GFTs and the DTL event intensity parameters.

When optimizing the GFTs (Step 1 in Fig. 3), GeneRax processes each family independently, and applies a tree search heuristic to each of them separately: for a given tree, it tests *all* possible *Subtree Prune and Regraft* (SPR) moves within a given radius and subsequently applies the SPR move that yields the tree with the highest joint likelihood. Then it iterates by again applying SPR moves to this new tree, until the joint likelihood can not be further improved. At the end of the GFT optimization, GeneRax increases the SPR radius by one until a certain maximum values is reached (see further below).

GeneRax optimizes the DTL intensity parameters globally over all gene families (Step 2 in Fig. 3). To this end, we apply the gradient descent method to find a set of DTL intensity parameters that maximizes the reconciliation likelihood over all gene families. We numerically approximate the gradient via finite differences.

The entire procedure stops when the SPR radius (starting from 1) exceeds a user-defined value. When the user does not define this maximum SPR radius, we set it to 5, as we did not observe any improvement above this value in our experiments.

### GFT and species tree reconciliation

The reconciliation likelihood computation algorithm conducts a post-order traversal of both, the species tree, and the GFT, and sums over all possible scenarios at each step of the traversal. To infer the ML reconciliation (Step 3 in Fig.3), GeneRax keeps track of the maximum likelihood path during the traversal.

GeneRax can export the reconciled GFTs into both Notung (Chen *et al*., 2000) and RecPhyloXML (Duchemin *et al*., 2018) formats (Fig. 4).

**FIG. 4.**
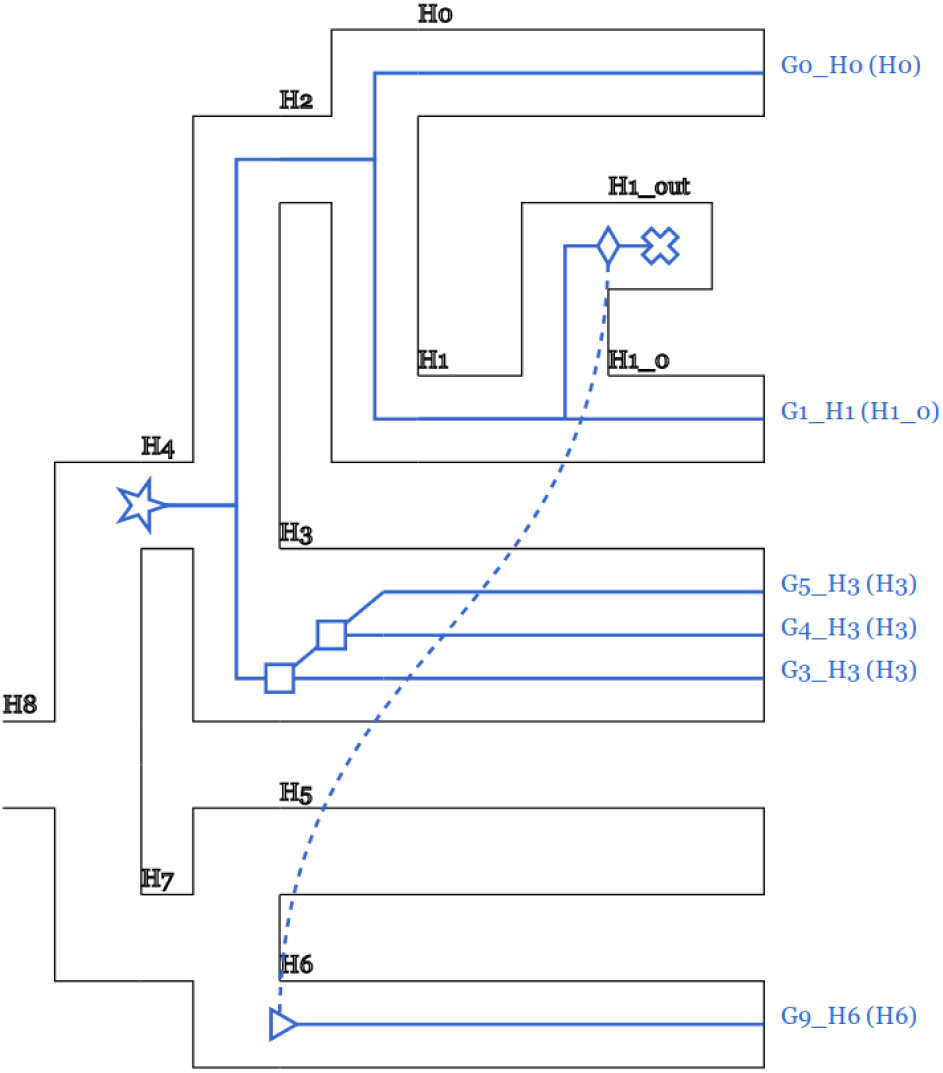
Reconciled GFT and species tree. Users can easily visualize reconciliations inferred with GeneRax using the online tool RecPhyloXML-visu (Duchemin *et al*., 2018). This example illustrates one HGT and one duplication events.

### Parallelization

Achieving ‘good’ parallel efficiency given a large number of gene families is challenging: the most straight-forward solution consists in assigning a subset of gene families to each core (Boussau *et al*., 2012). However, gene family MSAs are highly heterogeneous in terms of size, and are hence hard to evenly distribute over cores (Morel et al., 2018) such as to achieve ‘good’ load balance. In particular, large gene family MSAs can easily generate a parallel performance bottleneck. Our solution allows to split up individual inferences on such large gene family MSAs across several cores. Thus, we parallelize over, but also within gene families, in analogy to our ParGenes (Morel *et al*., 2018) tool. However, unlike ParGenes, GeneRax parallelizes individual GFT searches over the possible SPR moves and *not* over MSA sites. For a given GFT, we distribute the SPR moves we intend to apply among the cores assigned to the reconciliation of the GFT and apply them simultaneously. We adopted this parallelization approach for two reasons: (1) unlike the phylogenetic likelihood, the time for computing a reconciliation likelihood does not depend on the number of sites (i.e., a parallelization will not scale with the number of sites in contrast to the phylogenetic likelihood), and (2) per-MSA gene sequences are typically not long enough to efficiently parallelize the phylogenetic likelihood calculations over the sites.

## Experiments

We compared GeneRax to competing GFT inference methods on both, simulated, and empirical datasets.

### Tested software

This subsection describes the settings we used for executing the competing tools (summarized in Table 2) in all of our experiments.

We used ParGenes (Morel *et al*., 2018) to run RAxML-NG with 10 random and 10 parsimony starting trees and 100 bootstrap trees. For methods requiring starting GFTs, we selected the tree with the best likelihood found by RAxML-NG. We used 100 bootstrap trees to compute GFTs with branch support values as required for Notung and Treerecs. As Notung does not provide any explicit recommendation for setting the bootstrap support threshold, we used the default value (90%). We executed Treerecs with its automatic threshold selection from seven threshold values (seven is the default value). We executed Phyldog with a fixed species tree using a maximum SPR radius of 5, as in GeneRax, since Phyldog does not have a recommended setting. To execute ALE, we first generated posterior tree samples with MrBayes, using two independent runs, four chains, 1,000,000 generations, a sampling frequency of 1,000 and a burn-in of 100 trees. We used the undated ALE model to produce 100 tree samples per gene family. We used the same MrBayes tree samples to execute EcceTERA with the amalgamate option, without transfer from the dead, and with the dated species tree option.

Note that, Treerecs, Notung, MrBayes, EcceTERA, and ALE do not provide a parallelization over gene families for typical distributed memory compute cluster systems. To execute them on large datasets, we scheduled them with a dedicated MPI program, by dynamically assigning jobs (with one job per gene family) to the available MPI ranks, starting from the most expensive jobs with the largest gene family MSAs. Henceforth, we refer to *sequential runtime* as the sum of the time required by each program, and to *parallel runtime* as the elapsed time spent for the entire MPI run. For a given number of cores, the *parallel efficiency* is the sequential runtime divided by the product of the parallel runtime and the number of cores.

We executed GeneRax with default parameters and with both, random (GeneRax-random), and RAxML-NG (GeneRax-raxml) starting trees. When not stated otherwise, we present GeneRax results for random starting trees.

When working on simulated datasets that were not expected to contain HGT, we executed both, ALE, and GeneRax with a HGT rate set to zero, and denote these runs as ALE-DL and GeneRax-DL. When accounting for HGT, we denote them as ALE-DTL and GeneRax-DTL.

### Simulated datasets

We executed all tools listed in Table 2 on the dataset originally used to benchmark ALE (Szöllősi *et al*., 2013a). Szöllősi et al. initially inferred GFTs for 1099 Cyanobacteria gene families using ALE. Then, they simulated new sequences under the LG+Γ+I model along these trees, retaining both, the MSA sizes, and branch lengths. In our experiments, we inferred GFTs once under LG+Γ+I (true substitution model) and once under WAG without rate heterogeneity (misspecified substitution model).

In addition, we generated additional simulated datasets to investigate the influence of various parameters on the methods and their respective accuracy. The parameters we studied are the number of sites, the average gene branch lengths, the species tree size, and the DTL intensity parameters. We also used putative species trees that were increasingly different from the true species tree to quantify the robustness of the methods with respect to topological errors in the species tree. We simulated the species tree and GFTs using GenPhyloData (Sjöstrand *et al*., 2013), and the sequences using Seq-Gen (Rambaut and Grass, 1997), which simulates a continuous time birth and death process along a time-like species tree.

Finally, we executed simulations using SimPhy (Mallo *et al*., 2015) with increasing population sizes to assess the impact of ILS. We define the *ILS discordance* of a simulated dataset as being the average relative Robinson-Foulds (RF) distance (Robinson and Foulds, 1981) between the true species tree and the true GFTs obtained when running the same simulations without D, T, or L events.

### Empirical datasets

We executed all methods in Table 2 on the empirical datasets listed in Table 1. We measured both, sequential, and parallel runtimes. We also used GeneRax to evaluate the joint likelihood of the trees inferred with each method, to assess the quality of our tree search algorithm whose goal is to maximize this likelihood.

**Table 1.** Description of the empirical datasets used in our benchmarks. We extracted the Primates dataset from the release 96 of the Ensembl Compara database (Zerbino *et al*., 2017). The Cyanobacteria dataset was originally used in a previous study (Szöllősi *et al*., 2013a) and was extracted from the HOGENOM database (Penel *et al*., 2009).

| Dataset | Database | Species | Families | Avg. sites | Avg. genes | Max. genes |
| --- | --- | --- | --- | --- | --- | --- |
| Primates | ENSEMBL | 13 | 1523 | 84 | 45 | 349 |
| Cyanobacteria | HOGENOM | 36 | 1099 | 239 | 37 | 130 |

**Table 2.** Softwares used in our benchmark, with the type of method (ML, parsimony or both), the nature of the input trees (random tree, ML tree, tree with bootstrap support values or MCMC sample of trees), whether the method is STA and whether the method accounts for HGT.

| Software | Method type | Input trees | STA | HGT | Ref. |
| --- | --- | --- | --- | --- | --- |
| RAxML-NG | ML | Random | No | No | (Kozlov *et al.*, 2019) |
| Notung | Parsimony | Supported ML | Yes | No | (Chen *et al.*, 2000) |
| Treerecs | Parsimony + ML | Supported ML | Yes | No | (Comte *et al.*, 2018) |
| Phyldog | ML | ML | Yes | No | (Boussau *et al.*, 2012) |
| EcceTERA | Parsimony | Supported ML or MCMC samples | Yes | Yes | (Scornavacca *et al.*, 2014) |
| ALE | ML | MCMC samples | Yes | Yes | (Szöllősi *et al.*, 2013a) |
| GeneRax | ML | Random or ML | Yes | Yes | (this paper) |

## Results

In the following, we present the results of our experiments. For all methods, we report GFT quality (measured by RF distance to the true trees on simulated datasets, and joint likelihood on empirical datasets) and computational efficiency (measured by sequential runtime and parallel efficiency). All data and all inferred trees are available at https://cme.h-its.org/exelixis/material/generax_data.tar.gz.

### RF distances to true trees

We show the relative RF distances between the 1099 simulated Cyanobacteria true GTRs and the respective inferred GTRs in Fig. 5. For methods that yield more than one potential GFT per gene family (ALE and RAxML-NG), we average the distance over all inferred trees.

**FIG. 5.**
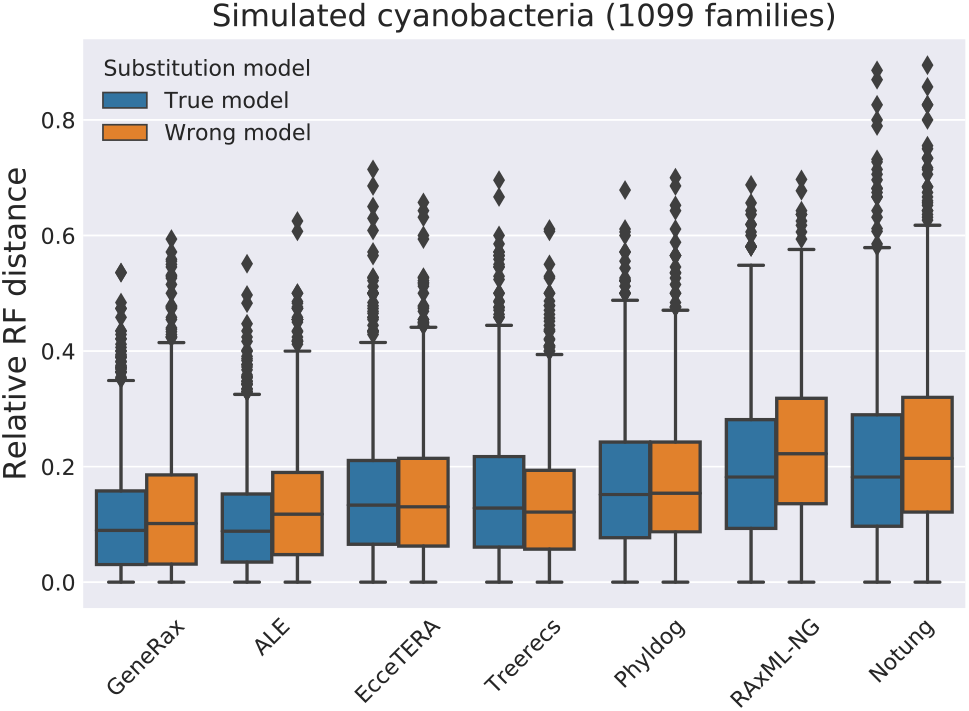
Relative RF distances to true trees, by inferring gene trees with the true substitution model (LG+Γ+I) and a misspecified substitution model (WAG).

GeneRax and ALE perform better than all other methods, except in the case of the misspecified substitution model where Treerecs performs equally well. Under the true model, STA methods that do not account for HGT but use a joint likelihood score (Phyldog and Treerecs) perform better than the purely sequence-based method (RAxML-NG), but worse than methods accounting for HGT. Although EcceTERA accounts for transfers, it only performs as good as Treerecs, presumably because the EcceTERA algorithm only uses parsimony. We hypothesize that Notung performs worse than all the other methods because it rearranges trees based on a parsimony score and an arbitrary support value threshold.

We summarize the results of the GenPhyloData simulations where we vary parameters (DTL intensity parameters, etc.) in presence of HGT in Fig. 6, and the results of the simulations in absence of HGT in the Supplementary Material. GeneRax finds the best trees in 90% of our simulation scenarios, but ALE finds trees that are almost as good in most simulations. Treerecs and Phyldog perform almost as well as GeneRax and ALE in the absence of HGT, but worse under HGT. Notung performs significantly worse than all SPA methods.

**FIG. 6.**
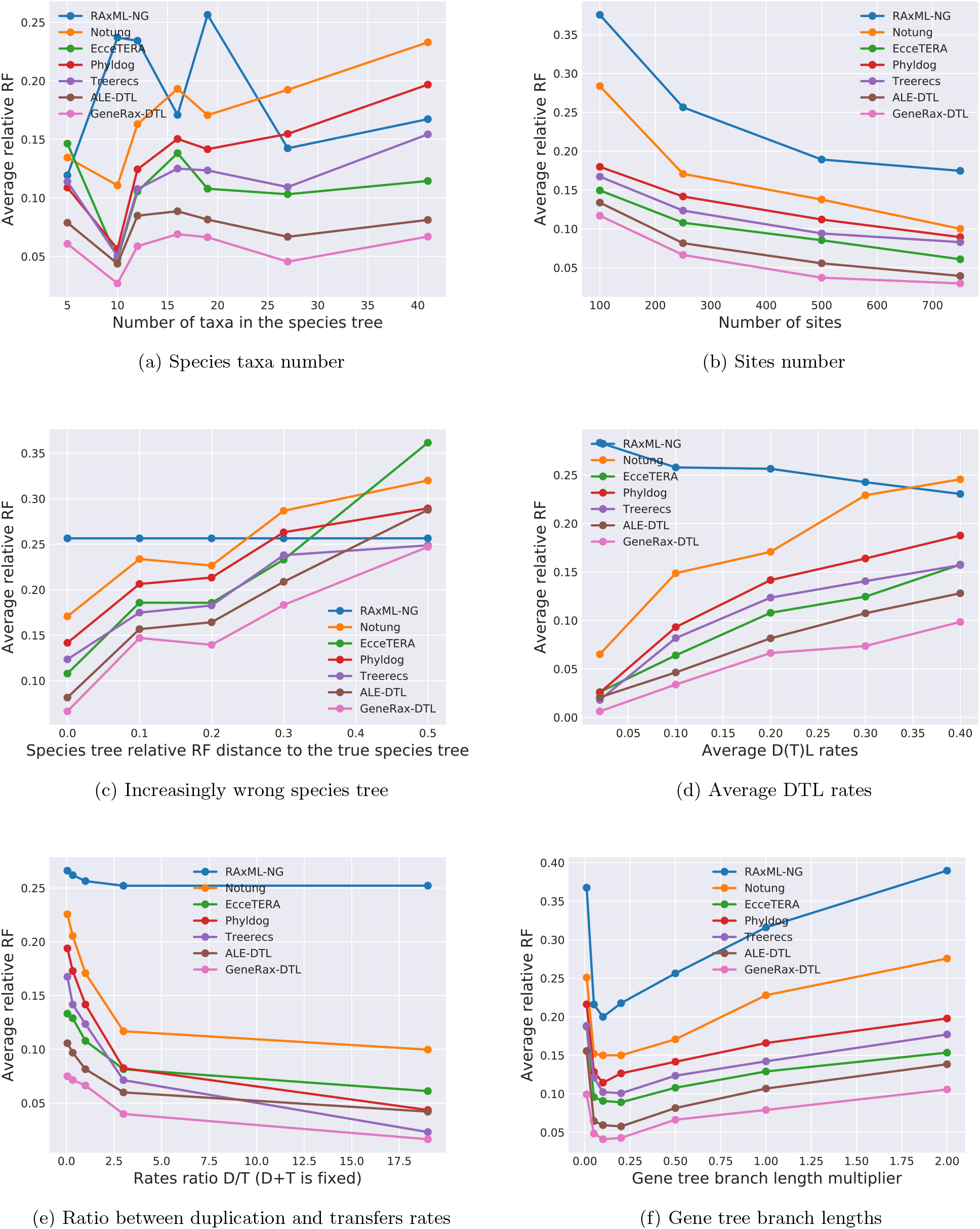
Comparison of different GTF correction tools on simulated datasets, in presence of horizontal gene transfers.

All STA methods show an analogous accuracy pattern when we vary parameters: they perform better with increasing gene sequence signal strength (Fig. 6 (b) and (f), and perform worse with increasing discordance between the species tree and the GFTs (Fig. 6 (c), (d) and (e)).

We show the results of the SimPhy simulations over varying ILS discordance scores in Fig. 7. GeneRax outperforms all other STA tools. It finds better GFTs than the only non-STA method (RAxML-NG) up to an ILS discordance score of 0.6. Our findings suggest that GeneRax can be deployed for analyzing datasets that exhibit a moderate degree of ILS.

**FIG. 7.**
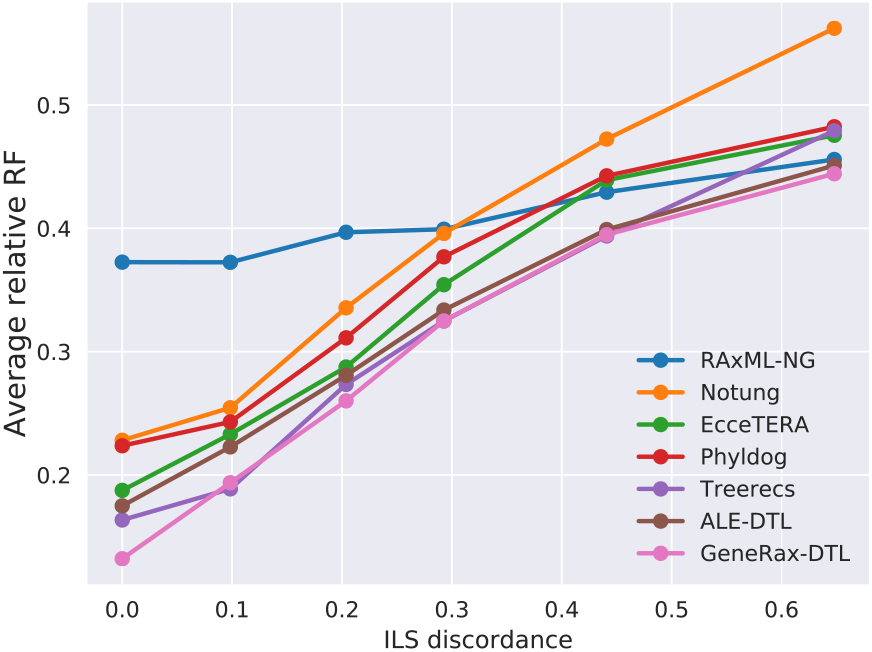
RF distance to true trees on simulated datasets with increasing discordance due to ILS.

### Branch score distances to true trees

To compare the quality of the gene branch lengths in terms of expected number of substitutions per site, we measured the average branch score distance (Kuhner and Felsenstein, 1994) between the inferred trees and the true trees (Fig. (8) with the phangorn R library (Schliep, 2010). GeneRax performs better than all competing tools. In particular, GeneRax shows a better average branch score distance (1.02) than ALE (1.48). A possible explanation for this is that ALE does not infer the branch lengths by optimizing the phylogenetic likelihood score, as opposed to GeneRax, Treerecs, and RAxML-NG. When using ALE, Notung, Phyldog, or EcceTERA, users interested in branch length accuracy would need to include an additional tool into their pipeline (e.g., RAxML-NG).

**FIG. 8.**
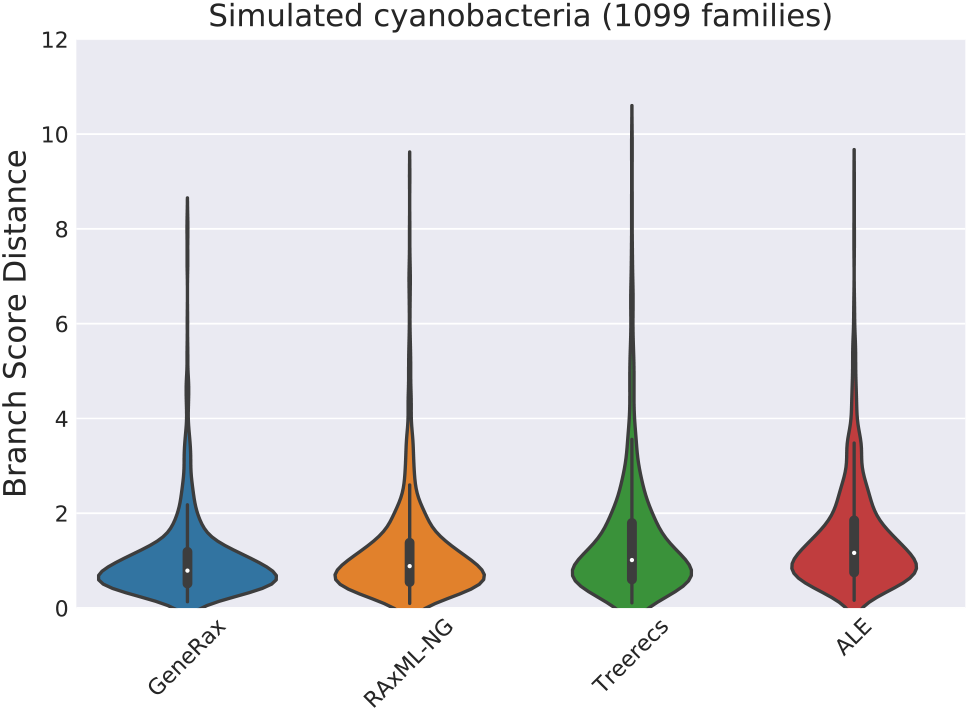
Branch score distance to true trees. We excluded from the plot methods that do not infer the branch lengths.

### Joint likelihood

We report the joint maximum likelihood scores of the GFTs obtained with the different tools in Fig. 9. As the true tree is generally not know for empirical data, and given that we are willing to accept the maximum likelihood criterion, we must assume that the tree yielding the best joint maximum likelihood is also the one that best explains the data. This approach of benchmarking ML tools on empirical datasets has been used repeatedly for assessing standard tree inference tools (Kozlov *et al*., 2019; Nguyen *et al*., 2015). The rationale for this is that standard tree searches based on the phylogenetic likelihood are inherently more difficult on empirical than on smooth and perfect simulated data. That is, differences between tree search algorithms might sometimes only be observable on empirical data. As expected, GeneRax finds the highest joint likelihood score. ALE is close to GeneRax, because it strives to approximate the same model. As the remaining tools implement distinct models, our comparison might appear as being unfair. However, we mainly regard this as a means of verifying that GeneRax properly maximizes the likelihood under its specific reconciliation model. Treerecs, Phyldog are also very close to GeneRax in absence of transfers, because they deploy a similar joint likelihood model. ALE performs better than Treerecs and Phyldog in presence of HGT, because Treerecs and Phyldog only account for gene duplication and loss. RAxML-NG, EcceTERA, and Notung do not implement a joint reconciliation likelihood model, which explains their low scores.

**FIG. 9.**
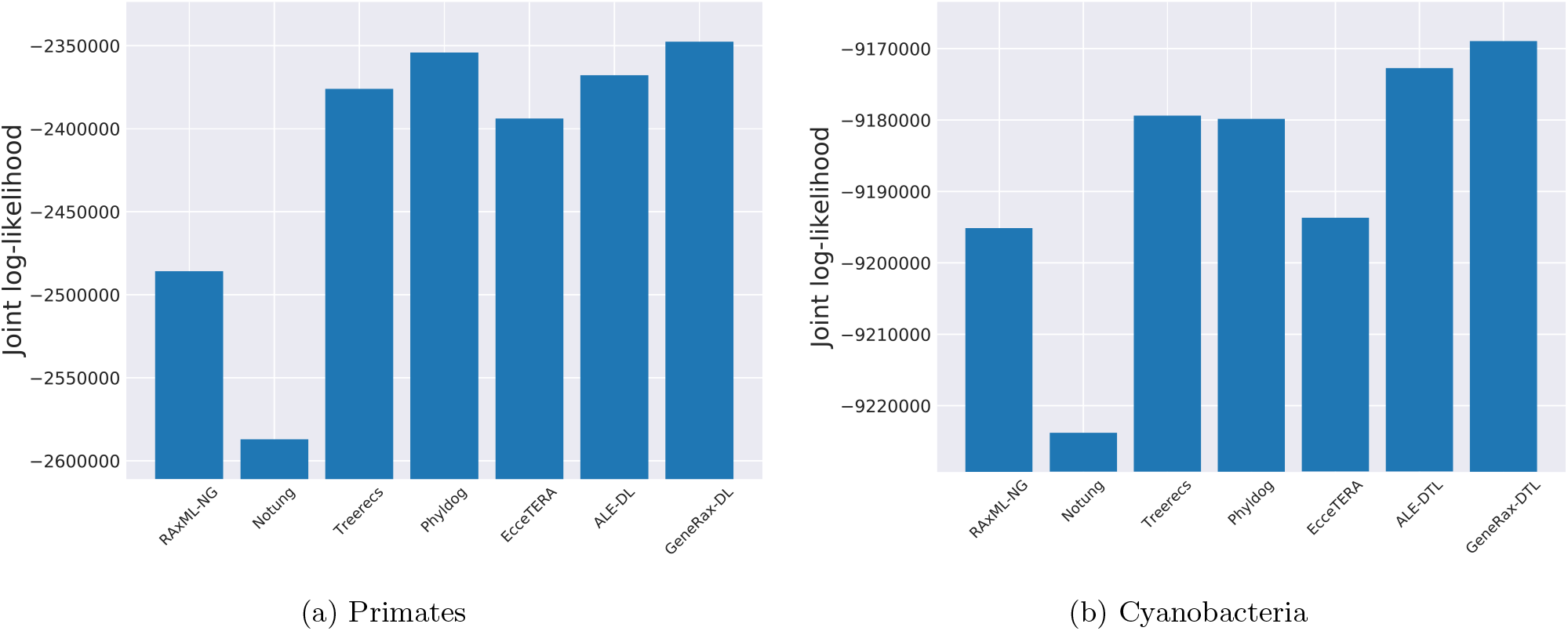
Log-likelihoods (the higher the better) evaluated with GeneRax. When evaluating the joint likelihood for Primates, we set the HGT rate to 0.

In addition, when running GeneRax on the empirical Cyanobacteria dataset, we recorded both, the reconciliation likelihood and the phylogenetic likelihood during the tree search (Fig. 10). We observe that the joint likelihood optimization occurs through an increase of the reconciliation likelihood in conjunction with a decrease of the phylogenetic likelihood. We observed this consistently on all simulated and empirical datasets we experimented with. In general, we observed that our joint likelihood tree search heuristic is not efficient in improving the phylogenetic likelihood score, and thus needs to start from trees with a high phylogenetic likelihood. For this reason, when the user does not provide a starting tree, we initially only optimize the phylogenetic likelihood, and only subsequently start the joint likelihood optimization.

**FIG. 10.**
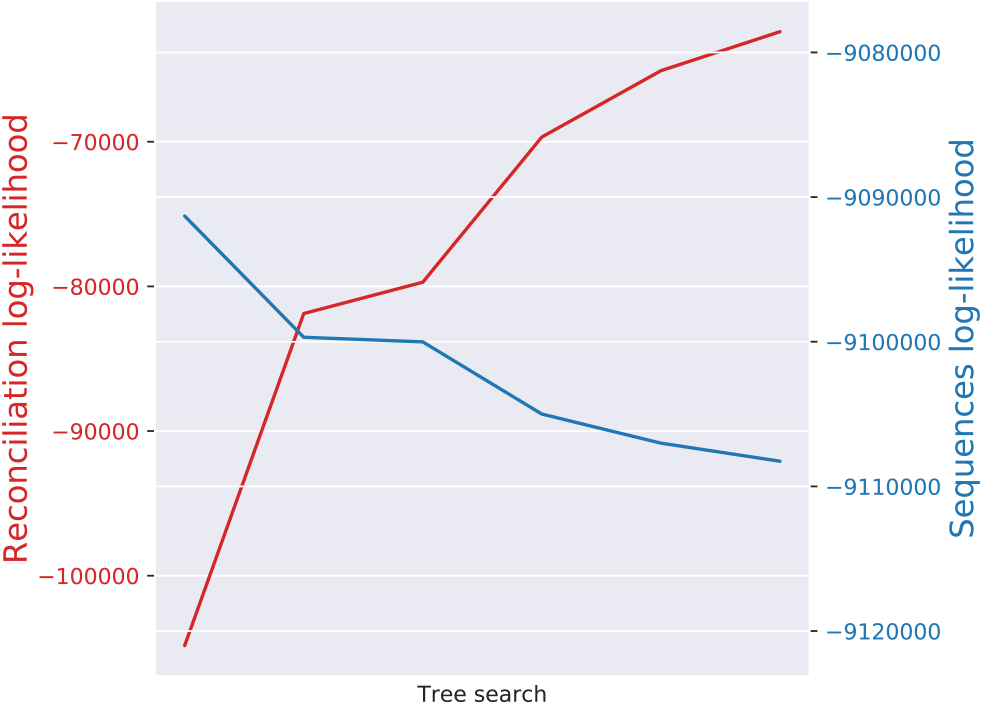
Reconciliation and sequence log-likelihoods during GeneRax tree search on the Cyanobacteria dataset. The sequence likelihood decreases while the reconciliation likelihood increases.

### Sequential runtimes

We measured the sequential runtimes of all tools on the empirical Cyanobacteria dataset. Comparing runtimes is not straightforward: some tools are very fast, but require an external preprocessing step, as described in Table 2. For instance, Notung is the fastest tool, but it requires GFTs with support values as input, and obtaining those can be extremely time-consuming. For a fair comparison, we plot both the time spent in the GFT inference tools alone, and the time spent in their respective pre-processing steps (Fig. 11).

**FIG. 11.**
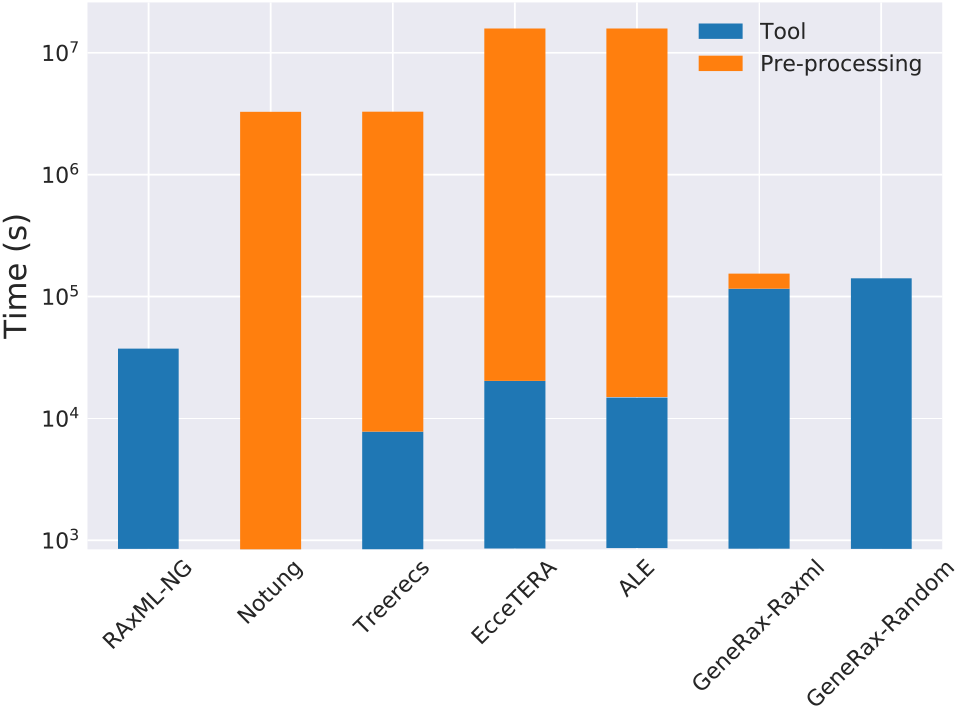
Sequential runtimes and additional overhead from precomputation steps (bootstrap trees with RAxML-NG for Notung and Treerecs, MCMC samples with MrBayes for ALE and EcceTERA, and RAxML-NG starting trees for GeneRax-raxml). The RAxML-NG column corresponds to the time spent in one single tree search. We represent times with a logarithmic scale.

When only considering the stand-alone runtimes of the tools, GeneRax is the slowest method. However, when including the pre-processing cost, GeneRax becomes the fastest STA approach. In addition, using only a single tool for the entire inference process substantially improves usability and reproducibility of the analyses.

### Parallel efficiency

We measured the parallel runtimes of GeneRax for different numbers of cores. For this experiment, we executed GeneRax on the empirical Cyanobacteria dataset (1099 families), starting from RAxML-NG trees. We used 4 up to 512 cores. Despite the highly heterogeneous gene family MSA sizes (in terms of both number of sites and number of taxa, see Supplementary Material), GeneRax achieves a high parallel efficiency of 70% on 512 cores. We plot the speedup as a function of the number of cores in the supplementary material.

We also measured the parallel efficiency of running the competing methods as described in the Experiments section, and plot them in the supplementary material. GeneRax is the only tool that achieves good efficiency (70%) because it parallelizes both, over, *and* within gene families, thereby achieving a ‘good’ load balance. Despite a similar two-level parallelization scheme, the parallel efficiency of RAxML-NG (scheduled with ParGenes, with one starting tree per family) is below 20%. The reason for this is that ParGenes parallelizes individual tree searches over the sites whereas GeneRax parallelizes them over the SPR moves. Gene MSAs are often short, and there is typically not a sufficient number of sites to allocate several cores per tree search with RAxML-NG. Other competing tools also fail to attain good parallel efficiency (40%), because they do not parallelize individual GFT inferences, and are thus limited by the longest individual per-tree inference time. The parallel efficiency of GeneRax decreases when starting from random trees, because the initial phylogenetic likelihood optimization step is based on RAxML-NG code, which does not implement our aforementioned two-level parallelization scheme yet.

## Discussion

### An accurate, robust and fast approach

We present GeneRax, an open source STA GFT inference software. GeneRax can simultaneously account for substitution and DTL events. It performs a tree search to optimize a joint likelihood, that is, the product of the phylogenetic likelihood and the reconciliation likelihood. It can handle multiple gene families in parallel. To the best of our knowledge, GeneRax is the first STA tool that does not require any pre-processing of the MSAs. Also, it does not require any arbitrary threshold settings or parsimony weights, and it can account for HGT.

On simulated datasets, we demonstrate that GeneRax and ALE find trees that are closer to the true trees than those inferred by competing tools. We show that GeneRax can provide more accurate gene family trees even when the species tree is inaccurate and the substitution model is misspecified. Using two empirical datasets (Cyanobacteria and Primates), we confirm that GeneRax finds the best-scoring maximum likelihood trees under its specific model among the tested tools, both, with, and without HGT. Finally, we show that GeneRax is not only faster than the tested competing methods (when accounting for the computational cost of the preprocessing steps), but also has a substantially higher parallel efficiency, making it suitable for seamless large-scale analyses.

GeneRax is a production-level code. We released it on BioConda (Grüning *et al*., 2018) to facilitate installation, and we kept its interface as simple as possible. While most competing STA methods require input GFTs, sometimes, including additional information (e.g., support values), GeneRax can directly infer the GFTs from a set of given MSAs. This simplified analysis process reduces the number of *ad hoc* choices that users have to make: GeneRax does not require bootstrap-support thresholds, parsimony weights, MCMC convergence criteria, chain settings, proposal tuning, or priors. Reducing the number of arbitrary choices does not only yield the tool easier to run, but also substantially improves the reproducibility of the results. One could contest the parameters we used in our experiments for the pre-processing steps: Treerecs and Notung *might* not need 100 bootstrap trees to obtain reliable support values. ALE and EcceTERA *might* not need as many MrBayes runs, chains, or generations to correctly approximate the phylogenetic likelihood. In general, it is possible to run the pre-processing steps faster than in our experiments. When running the competing methods, we tried to use the parameters that favor result quality/confidence over short runtimes, as we would have done in a real analysis.

### Limitations of GeneRax

GeneRax relies on two important assumptions: first, that the rooted species tree is known, and second, that the observed discordance between the GFTs and the species tree is mainly due to D, T, and L events. Our experiments suggest that, when those assumptions are violated, GeneRax can only improve the quality of the GFTs up to a certain degree. In particular, users should be cautious when using GeneRax in the presence of ILS. Furthermore, GeneRax is not suitable for improving GFT topologies in the presence of hybridization. Nonetheless, GeneRax might be deployed for detecting potential hybridization events, by identifying species pairs exhibiting an “abnormally high” number of HGT events.

### Future work

Despite the favorable evaluation results, GeneRax still faces several challenges.

First, the GeneRax reconciliation model does not take into account the branch lengths, neither in the species tree, nor in the GFTs. This leads to information loss, and furthermore allows for transfers between non-contemporary species. We believe that further adapting and extending the reconciliation model could improve the quality of the results. For instance, one could exploit an ultrametric dated species tree and use speciation events to slice the species tree, as done in (Szöllősi *et al*., 2012). However, slicing the species tree increases the number of inner species nodes quadratically, and thus incurs a substantial increase in computational cost.

Second, the GeneRax reconciliation model assumes that ILS does not occur. Some promising work (Chan *et al*., 2017; D Rasmussen and Kellis, 2012) has been conducted to combine DTL events and ILS in a single model. We believe that a computationally efficient software that can account for ILS, DTL events, and substitutions in a probabilistic framework would represent a major breakthrough in phylogenetic inference.

Finally, GeneRax needs a known/given species tree to estimate the GFTs. To this end, we plan to extend GeneRax to co-estimate both, the GFTs, and the species tree, as done in (Boussau *et al*., 2012). An approach to solving this challenge consists in inferring initial GFTs with non-STA methods, and then inferring an initial species tree that maximizes the reconciliation likelihood given these GFTs. Then, in a second step, one can propose new species tree topologies, optimize the GFTs and DTL intensity parameters on the proposed new species tree toplplogy, and update the species tree if the joint likelihood improves.

## Acknowledgments

This work was financially supported by the Klaus Tschira Foundation and by DFG grant STA 860/4-2. G.J.Sz. received funding from the European Research Council under the European Unions Horizon 2020 research and innovation programme under grant agreement no. 714774 and the grant GINOP-2.3.2.-15-2016-00057. We thank Bastien Boussau, Eric Tannier, Celine Scornavacca and Wandrille Duchemin for discussions on the topic. We also thank Laurent Duret and Simon Penel for their valuable user feedback.

## References

Åkerborg, Ö., Sennblad, B., Arvestad, L., and Lagergren, J. 2009. Simultaneous bayesian gene tree reconstruction and reconciliation analysis. Proceedings of the National Academy of Sciences, 106(14): 5714–5719.

Boussau, B., Szöllősi, G. J., Duret, L., Gouy, M., Tannier, E., Daubin, V., Lyon, U. D., and Lyon, U. 2012. Genome-scale coestimation of species and gene trees. Life Sciences, pages 1–27.

Chan, Y., Ranwez, V., and Scornavacca, C. 2017. Inferring incomplete lineage sorting, duplications, transfers and losses with reconciliations. Journal of Theoretical Biology, 432: 1–13.

Chen, K., Durand, D., and Farach-Colton, M. 2000. Notung: A program for dating gene duplications and optimizing gene family trees. Journal of computational biology: a journal of computational molecular cell biology, 7: 429–47.

Comte, N., Morel, B., Hasic, D., Guéguen, L., Boussau, B., Daubin, V., Scornavacca, C., Gouy, M., Stamatakis, A., Tannier, E., and Parsons, D. 2018. Treerecs. https://gitlab.inria.fr/Phylophile/Treerecs/tree/pll-integration.

D Rasmussen, M. and Kellis, M. 2012. Unified modeling of gene duplication, loss, and coalescence using a locus tree. Genome research, 22: 755–65.

Darriba, D., Flouri, T., Kozlov, A., Morel, B., and Stamatakis, A. 2019. Pll-modules.

Duchemin, W., Gence, G., Arigon Chifolleau, A.-M., Arvestad, L., Bansal, M. S., Berry, V., Boussau, B., Chevenet, F., Comte, N., Davn, A. A., Dessimoz, C., Dylus, D., Hasic, D., Mallo, D., Planel, R., Posada, D., Scornavacca, C., Szöllősi, G., Zhang, L., Tannier,., and Daubin, V. 2018. RecPhyloXML: a format for reconciled gene trees. Bioinformatics, 34(21): 3646–3652.

El-Mabrouk, N. and Noutahi, E. 2019. Gene Family EvolutionAn Algorithmic Framework, pages 87–119. Springer International Publishing.

Felsenstein, J. 1981. Evolutionary trees from dna sequences: A maximum likelihood approach. Journal of Molecular Evolution, 17(6): 368–376.

Felsenstein, J. 1985. Confidence limits on phylogenies: an approach using the bootstrap. Evolution, 39(4): 783–791.

Grüning, B., Dale, R., Sjödin, A., Chapman, B., Rowe, J., Tomkins-Tinch, C., Valieris, R., Köster, J., Blin, K., Haudgaard, M., Kratz, A., Junge, A., and Knudsen, M. 2018. Bioconda: sustainable and comprehensive software distribution for the life sciences. Nature Methods, 15: 475–476.

Kozlov, A. M., Darriba, D., Flouri, T., Morel, B., and Stamatakis, A. 2019. RAxML-NG: a fast, scalable and user-friendly tool for maximum likelihood phylogenetic inference. Bioinformatics.

Kuhner, M. K. and Felsenstein, J. 1994. A simulation comparison of phylogeny algorithms under equal and unequal evolutionary rates. Molecular Biology and Evolution, 11(3): 459–468.

Larget, B. 2013. The estimation of tree posterior probabilities using conditional clade probability distributions. Systematic biology, 62.

Mallo, D., De Oliveira Martins, L., and Posada, D. 2015. SimPhy: Phylogenomic Simulation of Gene, Locus, and Species Trees. Systematic Biology, 65(2): 334–344.

Mirarab, S., Reaz, R., Bayzid, M., Zimmermann, T., S Swenson, M., and Warnow, T. 2014. Astral: Genome-scale coalescent-based species tree estimation. Bioinformatics (Oxford, England), 30: i541–i548.

Morel, B., Kozlov, A. M., and Stamatakis, A. 2018. ParGenes: a tool for massively parallel model selection and phylogenetic tree inference on thousands of genes. Bioinformatics.

Musilova, Z., Cortesi, F., Matschiner, M., Davies, W. I. L., Patel, J. S., Stieb, S. M., de Busserolles, F., Malmstrøm, M., Tørresen, O. K., Brown, C. J., Mountford, J. K., Hanel, R., Stenkamp, D. L., Jakobsen, K. S., Carleton, K. L., Jentoft, S., Marshall, J., and Salzburger, W. 2019. Vision using multiple distinct rod opsins in deep-sea fishes. Science, 364(6440): 588–592.

Nguyen, L.-T., Schmidt, H. A., von Haeseler, A., and Minh, B. Q. 2015. IQ-TREE: A Fast and Effective Stochastic Algorithm for Estimating Maximum-Likelihood Phylogenies. Molecular Biology and Evolution, 32(1): 268–274.

Noutahi, E., Semeria, M., Lafond, M., Seguin, J., Boussau, B., Guguen, L., El-Mabrouk, N., and Tannier, E. 2016. Efficient gene tree correction guided by genome evolution. PLOS ONE, 11.

Penel, S., Arigon, A.-M., Dufayard, J.-F., Sertier, A.-S., Daubin, V., Duret, L., Gouy, M., and Perrière, G. 2009. Databases of homologous gene families for comparative genomics. BMC Bioinformatics, 10(6): S3.

Rambaut, A. and Grass, N. C. 1997. Seq-Gen: an application for the Monte Carlo simulation of DNA sequence evolution along phylogenetic trees. Bioinformatics, 13(3): 235–238.

Robinson, D. and Foulds, L. 1981. Comparison of phylogenetic trees. Mathematical Biosciences, 53(1): 131–147.

Schliep, K. P. 2010. phangorn: phylogenetic analysis in R. Bioinformatics, 27(4): 592–593.

Scornavacca, C., Jacox, E., and Szöllősi, G. J. 2014. Joint amalgamation of most parsimonious reconciled gene trees. Bioinformatics, 31(6): 841–848.

Sennblad, B. and Lagergren, J. 2009. Probabilistic Orthology Analysis. Systematic Biology, 58(4): 411–424.

Sjöstrand, J., Arvestad, L., Lagergren, J., and Sennblad, B. 2013. Genphylodata: realistic simulation of gene family evolution. BMC Bioinformatics, 14(1): 209.

Szöllősi, G. J., Boussau, B., Abby, S. S., Tannier, E., and Daubin, V. 2012. Phylogenetic modeling of lateral gene transfer reconstructs the pattern and relative timing of speciations. Proceedings of the National Academy of Sciences, 109(43): 17513–17518.

Szöllősi, G. J., Rosikiewicz, W., Boussau, B., Tannier, E., and Daubin, V. 2013a. Efficient exploration of the space of reconciled gene trees. Systematic Biology, 62(6): 901–912.

Szöllősi, G. J., Tannier, E., Lartillot, N., and Daubin, V. 2013b. Lateral Gene Transfer from the Dead. Systematic Biology, 62(3): 386–397.

Szöllősi, G. J., Tannier, E., Daubin, V., and Boussau, B. 2014. The Inference of Gene Trees with Species Trees. Systematic Biology, 64(1): e42–e62.

Touchon, M., Hoede, C., Tenaillon, O., Barbe, V., Baeriswyl, S., Bidet, P., Bingen, E., Bonacorsi, S., Bouchier, C., Bouvet, O., Calteau, A., Chiapello, H., Clermont, O., Cruveiller, S., Danchin, A., Mdric, D., Dossat, C., El Karoui, M., Frapy, E., and Denamur, E. 2009. Organised genome dynamics in the escherichia coli species results in highly diverse adaptive paths. PLoS genetics, 5: e1000344.

Yang, Z. 1994. Statistical properties of the maximum likelihood method of phylogenetic estimation and comparison with distance matrix methods. Systematic Biology, 43(3): 329–342.

Zerbino, D. R., Achuthan, P., Akanni, W., Amode, M., Barrell, D., Bhai, J., Billis, K., Cummins, C., Gall, A., Girn, C. G., Gil, L., Gordon, L., Haggerty, L., Haskell, E., Hourlier, T., Izuogu, O. G., Janacek, S. H., Juettemann, T., To, J. K., Laird, M. R., Lavidas, I., Liu, Z., Loveland, J. E., Maurel, T., McLaren, W., Moore, B., Mudge, J., Murphy, D. N., Newman, V., Nuhn, M., Ogeh, D., Ong, C. K., Parker, A., Patricio, M., Riat, H. S., Schuilenburg, H., Sheppard, D., Sparrow, H., Taylor, K., Thormann, A., Vullo, A., Walts, B., Zadissa, A., Frankish, A., Hunt, S. E., Kostadima, M., Langridge, N., Martin, F. J., Muffato, M., Perry, E., Ruffier, M., Staines, D. M., Trevanion, S. J., Aken, B. L., Cunningham, F., Yates, A., and Flicek, P. 2017. Ensembl 2018. Nucleic Acids Research, 46(D1): D754–D761.

Zheng, Y. and Zhang, L. 2014. Effect of incomplete lineage sorting on tree-reconciliation-based inference of gene duplication. IEEE/ACM Transactions on Computational Biology and Bioinformatics, 11(3): 477–485.

