## SupplementMaterial for "GeneRax: A tool for species tree-aware maximum likelihood based gene family tree inference under gene duplication, transfer, and loss"

### Abstract

Supplement material for GeneRax paper.

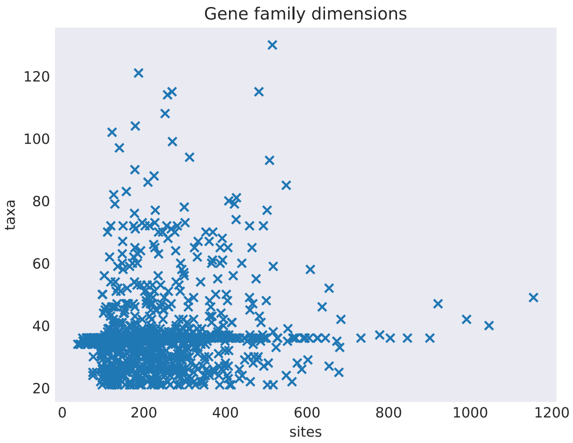


**FIG. 1.** Gene family dimension in the cyanobacteria dataset. Each point is a gene family. The number of sites is the number of unique sites (we do not count duplicates).

© The Author 2013. Published by Oxford University Press on behalf of the Society for Molecular Biology and Evolution. All rights reserved. For permissions, please

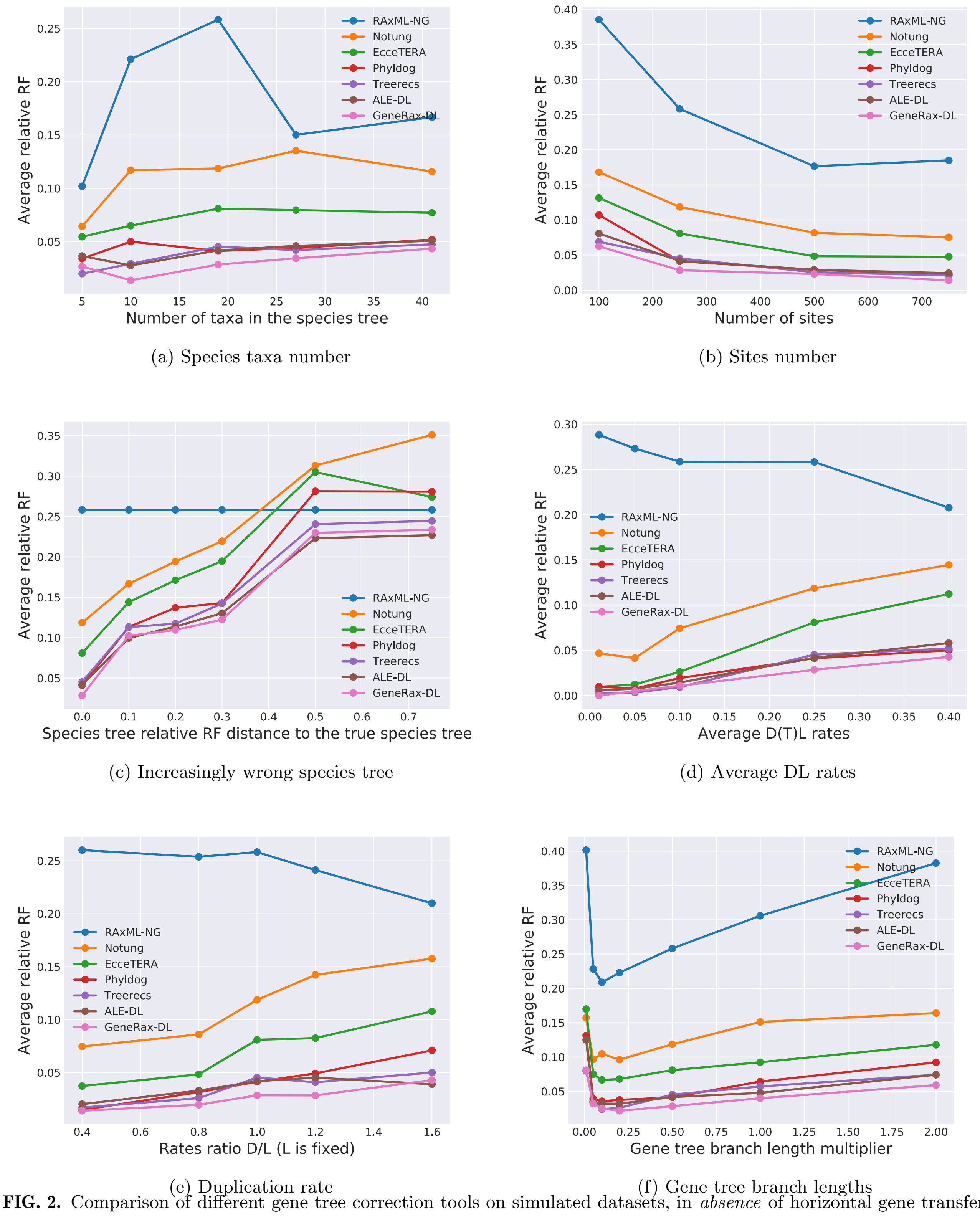


**FIG. 2.** Comparison of different gene tree correction tools on simulated datasets, in *absence* of horizontal gene transfers.

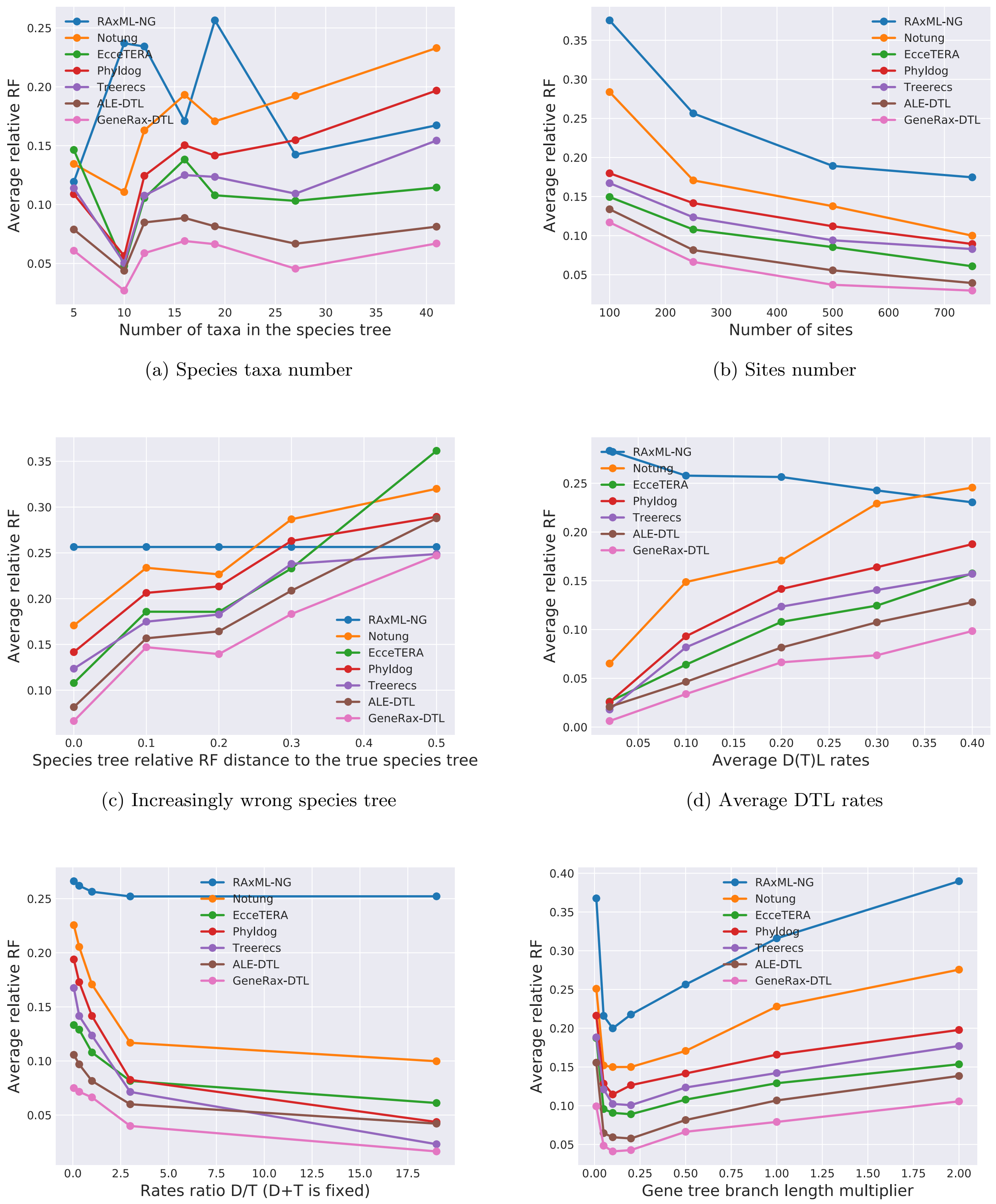


(a) Species taxa number (b) Sites number

(c) Increasingly wrong species tree (d) Average DTL rates

(e) Ratio between duplication and transfers rates (f) Gene tree branch lengths

**FIG. 3.** Comparison of different GTF correction tools on simulated datasets, in *presence* of horizontal gene transfers.

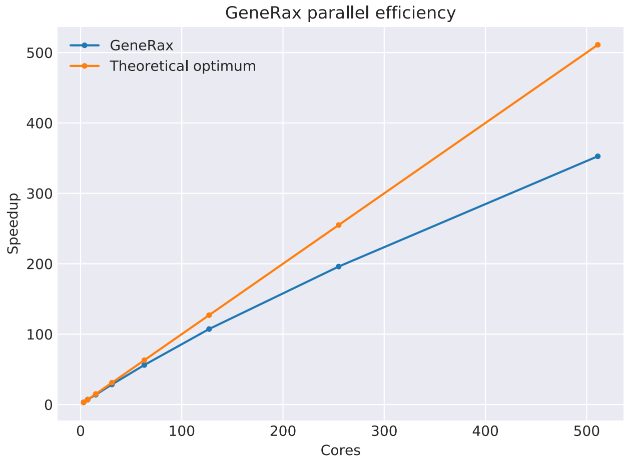


**FIG. 4.** Parallel speedup of GeneRax on the empirical Cyanobacteria dataset (1099 families), using from 4 to 512 cores. What about moving this to supplement?

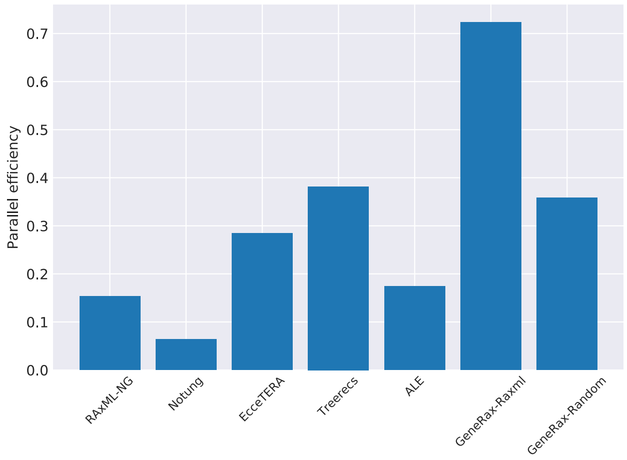


**FIG. 5.** Parallel efficiency of the different methods applied to the cyanobacteria empirical dataset on 512 CPU cores. We do not include pre-processing steps.
